# Intergenerational metabolic priming by sperm piRNAs

**DOI:** 10.1101/2021.03.29.436592

**Authors:** Adelheid Lempradl, Unn Kugelberg, Mary Iconomou, Ian Beddows, Daniel Nätt, Eduard Casas, Lovisa Örkenby, Lennart Enders, Alejandro Gutierrez Martinez, Oleh Lushchak, Erica Boonen, Tamina Rückert, Martin Sabev, Marie G.L. Roth, Dean Pettinga, Tanya Vavouri, Anita Öst, J. Andrew Pospisilik

## Abstract

**Summary:** Preconception parental environment can reproducibly program offspring phenotype without altering the DNA sequence, yet the mechanisms underpinning this ‘epigenetic inheritance’ remains elusive. Here, we demonstrate the existence of an intact piRNA-pathway in mature *Drosophila* sperm and show that pathway modulation alters offspring gene transcription in a sequence-specific manner. We map a dynamic small RNA content in developing sperm and find that the mature sperm carry a highly distinct small RNA cargo. By biochemical pulldown, we identify a small RNA subset bound directly to piwi protein. And, we show that piRNA-pathway controlled sperm small RNAs are linked to target gene repression in offspring. Critically, we find that full piRNA-pathway dosage is necessary for the intergenerational metabolic and transcriptional reprogramming events triggered by high paternal dietary sugar. These data provide a direct link between regulation of endogenous mature sperm small RNAs and transcriptional programming of complementary sequences in offspring. Thus, we identify a novel mediator of paternal intergenerational epigenetic inheritance.

## Introduction

Current data suggest obesity as one of the world’s chief socioeconomic challenges of our day, impacting ∼1 billion individuals worldwide. The dramatic rise in metabolic disease incidence in the last decades, particularly in children, suggests a prominent role for epigenetic mechanisms, in particular, a role for intergenerational epigenetic mechanisms where physiological effects in parents (e.g. diet, hyperglycemia, obesity) trigger disease-predisposing shifts in the offspring through non-DNA-sequence-based mechanisms. To date, the mechanisms mediating intergenerational epigenetic programming in response to physiological state remain poorly understood.

PIWI-interacting RNAs (piRNAs) are 24-32 nucleotide, PIWI-bound small RNAs that are best known for silencing transposable elements and thereby limiting their mutagenic potential^1-3^. This canonical function of piRNAs is essential for germline genome integrity and is active in most animals^4^. The piRNA-pathway differs from other small RNA (sRNA) pathways (miRNA, siRNA) in three key aspects: 1) piRNA-pathway protein expression is mainly restricted to reproductive organs; 2) piRNAs are generated through a Dicer-independent mechanism; and 3) piRNAs are processed from single stranded precursor transcripts, making mRNAs theoretical sources and substrates for piRNA generation and amplification.

Canonical piRNAs are derived from genome regions called piRNA clusters, which harbor ancient transposon fragments. Accumulating evidence in both flies and mice suggests that piRNAs can also be produced from genic mRNAs^5-10^. In male mice, about 20% of the piRNA population in pre-pachytene germ cells for example, is derived from the exons of hundreds of mRNAs^5^. piRNAs derived from 3’UTRs have also been found in follicle cells of fly ovaries and Xenopus eggs^6,7,10^. Although piRNAs were first identified in *Drosophila* testis^11^, much of the pioneering work in flies has focused on the female germline^12^. In general, the sRNA content of the male *Drosophila* germline is ill characterized. Specifically, piRNA populations, their dynamics during spermatogenesis and their potential functional roles, remain poorly understood.

In *C. elegans*, piRNAs have been implicated in the multigenerational inheritance of foreign DNA-triggered epigenetic silencing^13^. In these contexts, once piRNA-seeded silencing states are initially established, their long-term memory is independent of the original piRNA trigger, and instead relies on nuclear RNAi and chromatin pathways. Notable, maternal piRNAs have been shown to buffer against DNA sequence incompatibility between maternal and paternal genomes, in particular at transposons, a phenomenon known as hybrid dysgenesis^14^. Here, we identify an intact mature sperm piRNA pathway and provide evidence for its involvement in both encoding and decoding intergenerational inheritance effects.

### Dynamic sRNA expression in the male ***Drosophila*** germline

The tube-shaped *Drosophila* testis has contributed significantly to our understanding of stem cell maintenance and germ cell differentiation^15^. To understand the dynamics of sRNA populations during spermatogenesis we performed sRNA sequencing on testes manually dissected into four parts: 1) testis tip (T1) containing stem cells and primary spermatocytes, 2) the apical portion (T2) containing meiotic and developing spermatogonia, 3) the distal portion (T3) containing late-stage spermatocytes undergoing individualization, and 4) mature sperm (Sperm) isolated from seminal vesicles (Fig. 1A-C). Principal component analysis revealed high technical reproducibility. Interestingly, the greatest variation (PC1) separated sperm from testis samples; PC2 separated the three stages of testis development (Fig. 1B). These data indicate that the mature sperm sRNA repertoire is highly distinct from that of the developing germ cells. Relative to T1-T3, sperm showed an *increase* in sRNAs mapping to protein coding genes (3’ UTRs and exons) and tRNAs, and a *decrease* in repeat-associated sRNAs (piRNA clusters, complex and simple repeats) (Fig. 1C). These results resemble findings in the male mouse germline, where a loss of piRNAs and a gain of tRNA and mRNA fragments was shown for the transition from testicular to caput sperm^16^. Sperm sRNAs mapping to protein coding exons exhibited a high correlation with sperm mRNA-seq datasets (r=0.92, p<0.0001; Fig. 1D), consistent with the mRNA degradation that occurs in late spermatogenesis. Fly and human sperm^17^ sRNA (Fig. 1E and F) repertoires showed critical similarity at the biotype level, sharing for instance seven of the ten most highly expressed tRNA genes (Fig. 1G), thus highlighting the evolutionary conservation. These results comprise the first in-depth analysis of sRNA dynamics in *Drosophila* spermatogenesis; they demonstrate a highly specific sperm sRNA population relative to the developing germline; and, they show conservation in sRNA composition between *Drosophila* and mammals.

**Fig. 1:**
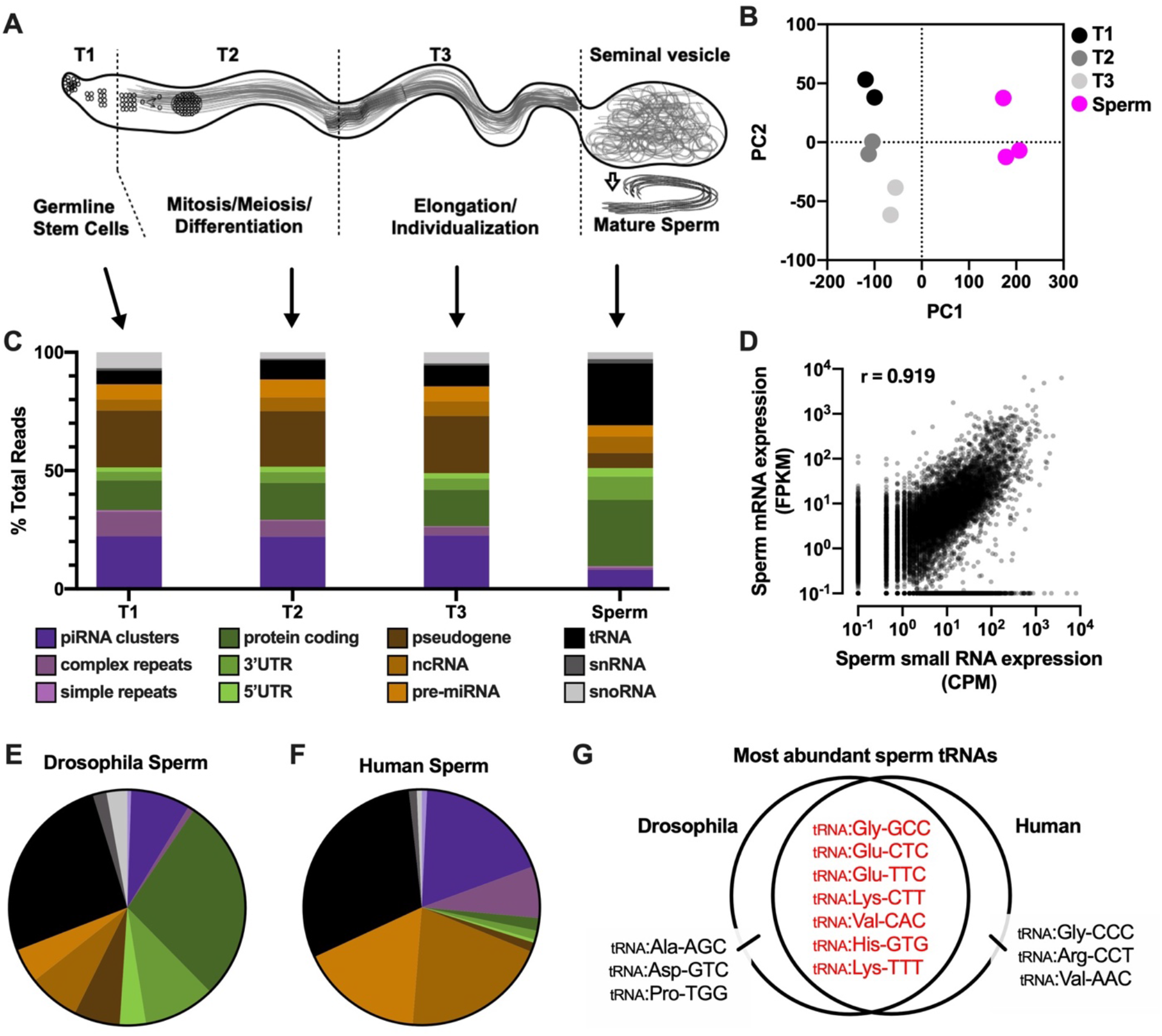
Dynamic sRNA expression in the male *Drosophila* germline. A) A schematic of the *Drosophila* testis indicating segments used for sRNA sequencing; B) PCA plot of sRNA sequencing data; C) % read distribution across biotypes and across testis segments; biotype color legend applies to panels C, E and F; D) correlation of sRNA sequencing results with previously published mRNA sequencing^30^ results from mature sperm and showing Peason’s r; E) *Drosophila* sperm and F) human sperm sRNA biotype distributions; G) 10 most enriched tRNA features in *Drosophila* and human sperm^17^ sRNA sequencing.

### Evidence for a functional piRNA-pathway in mature sperm

One key outstanding question about sperm sRNAs in the context of intergenerational control is how the minimal RNA load of a single sperm cell can reprogram the developmental trajectory of a whole organism. The amplification mechanism of the piRNA-pathway provides one possibility by which quantal differences in sRNAs might trigger reproducible next-generation effects as well as a range of penetrance distributions. piRNAs were originally identified and characterized based on their direct binding to PIWI proteins ^18-22^. If piRNAs were involved in epigenetic inheritance, we reasoned that they would be loaded onto the appropriate protein machinery in mature sperm. Previous work showed that Piwi protein is stably expressed in somatic and early germ-cells in the apical testis ^23^. Consistent with those reports, we found Piwi immunoreactivity in the nuclei of multiple germline and somatic cell types in the tip of the testis (Supplemental Fig. 1A). Interestingly, we also found piwi protein in individualizing, late stage spermatids at the distal end of the testis, and in washed and purified sperm isolated from the sperm sack when using anti-Piwi antibody raised against a peptide from the middle of the protein (Fig. 2A left; see Materials and Methods for antibody details;). Piwi immunoreactivity appeared both as puncta along the sperm tail (Fig. 2A center) and as a highly reproducible single body at the base of each compacted sperm nucleus (Fig. 2A right, and secondary antibody control in Supplemental Fig. 1B). This result is in agreement with results from mouse showing PiwiL1 protein presence in mature sperm^24^. Western blots confirmed our result by detecting the full-length Piwi protein in isolated sperm and ovaries (∼97 kDa; Western blot, Fig. 2B left and middle panel). Samples containing empty seminal vesicles showed no evidence of Piwi protein, which argues against possible contribution of somatic cell contamination to the signal in sperm. Further, we isolated protein lysates from ovaries, testis, and purified sperm of flies carrying a BAC Piwi-GFP transgene. In contrast to wild-type, Western blot of Piwi-GFP transgenic sperm revealed two bands, matching the sizes of GFP-tagged and wild-type Piwi protein (Fig. 2C). Thus, both wild-type and transgenic Piwi protein are found in sperm, confirming the specificity of the antibody.

**Fig. 2:**
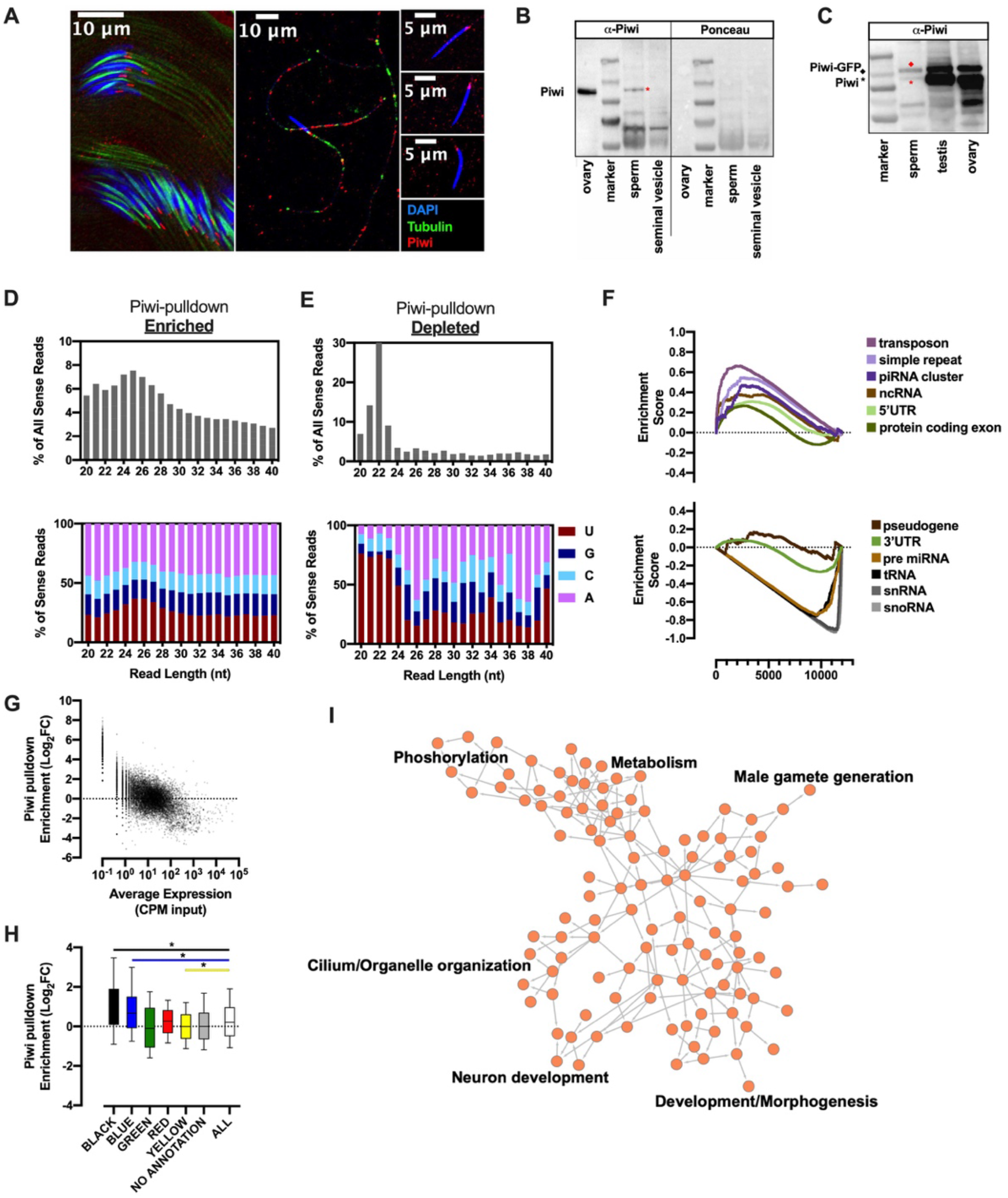
Presence of Piwi bound RNAs in mature *Drosophila* sperm. A) immunofluorescence staining of sperm with Piwi (ab5207) and tubulin antibodies; B) left panel shows western blot with Piwi antibody (ab5207) for ovary, sperm and empty seminal vesicle, right panel shows Ponceau staining to show protein content per lane. C) sperm, testis and ovaries from GFP-Piwi flies, the upper band shows the GFP fusion protein, lower band shows WT Piwi; D-E) length distribution (left) and 1^st^ nucleotide bias (right) of sense reads mapping to all features displayed as percentages of all reads between the length of 20-50nt for significantly enriched (D) and for significantly depleted features in Piwi-IP (piwi-antibody ab5207)(E); F) GSEA analysis of sRNA results using custom pathways for the individual biotypes used for annotation; G) average counts per million (CPM) in input samples versus Log_2_FC in Piwi pulldown samples H) enrichment of reads mapping to features embedded in different chromatin states. I) Pathway overrepresentation analysis (WebGestalt)^31^ of protein coding features significantly enriched in Piwi pulldown.

Seeking to leverage an antibody-independent technique, we performed targeted mass spectrometry on protein extracts of purified sperm (Supplemental Fig. 1C). For Piwi, we found signals for all possible targetable Piwi peptides after tryptic digestion. Importantly, peptides were also detected for the remaining two PIWI proteins known to be necessary for amplification of silencing competent piRNAs, namely aubergine (Aub) and argonaute 3 (AGO3)(Supplemental Fig. 1C). Thus, intact PIWI pathway proteins are consistently expressed in mature sperm and may therefore serve as facilitators of intergenerational inheritance of functional piRNAs.

To identify piwi-associated sRNAs we used the sperm specific Piwi antibody to develop and optimize a biochemical pulldown and sequencing approach (Piwi-RIP-seq) applicable to the minute amounts of Piwi protein and RNA available in dissected *Drosophila* sperm. After rounds of protocol optimization, we profiled the Piwi-bound sRNA content in sperm from 4 replicates each of ∼750 hand-dissected seminal vesicles (Piwi-RIP-seq). Piwi-pulldown enriched sRNAs showed a peak length of ∼25nt (Fig. 2D left) and an A-bias at the first nucleotide position (Fig. 2D right), consistent with published data from ovaries^18^. Using a stringent mapping and hierarchical annotation approach and rRNA exclusion (see methods), we found enrichment for sRNAs from piRNA clusters, repeats and transposons, and to a lesser extent those from protein coding exons, 5’UTRs and ncRNAs (Fig. 2F, top). By contrast, sRNAs *not* associated with piwi protein in the pull-down experiment showed a peak length of ∼21-23 nt (Fig. 2E) and were enriched for miRNAs, tRNAs, snRNAs, snoRNAs pseudogenes and 3’UTRs (Fig. 2F, bottom). Importantly, Piwi-bound RNAs exhibited an anti-correlation with total sRNA content of sperm (Fig. 2G). These data indicated highly specific loading of sRNAs onto Piwi and also argued against non-specific pulldown effects in the Piwi-RIP-seq results. Pathway analysis of protein coding exon-derived sRNAs showed enrichments for genes involved in phosphorylation, metabolism, gametogenesis, development and cilium organization (Fig. 2I). To test for potential chromatin state associated enrichment we next mapped our Piwi-RIP-seq dataset to published *Drosophila* chromatin state annotations derived from genome-wide binding analysis of over 50 functional chromatin-binding proteins^25^. Interestingly, this intersection showed a striking enrichment for sRNAs from transcripts that map to chromatin states with clear repressive signatures. These enriched states were described as Lamin / H1-associated, “Black” chromatin and Polycomb-associated, “Blue” chromatin in embryonic cells (Fig. 2H and Fig S1 D). Thus, mature sperm harbors a highly specific complement of Piwi-bound sRNAs (piRNAs).

### Piwi controlled sperm RNAs regulate offspring gene transcription

Working from the hypothesis that sperm piRNAs might regulate offspring transcription, we examined the effect of *parental* heterozygous *aub* mutation (*aub*^*Het*^) on next-generation transcriptional output (embryos, stage 17). Aub participates in the ping-pong cycle of the PIWI pathway and is necessary for amplification of silencing competent piRNAs and robust gene repression^26,27^. We compared mRNA transcriptional changes elicited in offspring from two parallel mutant crosses (Fig. 3A and B): *aub*^*Het*^ mothers x WT fathers (maternal *aub*^*Het*^ offspring); and WT mothers x *aub*^*Het*^ fathers (paternal *aub*^*Het*^ offspring)(data expressed relative to wildtype offspring from parallel WT x WT crosses; WT offspring). In order to ensure comparable genetic backgrounds, we prepared for the experiment by backcrossing *aub*^*Het*^ flies to our own highly inbred w1118 background (WT) (see supplemental methods for details). The mRNA transcriptional response in maternal *aub*^*Het*^ offspring revealed a strong correlation with our RIP-seq defined sperm piRNAs (Fig. 3C), specifically, derepression of piRNA-sequence matching ‘targets’. These data suggested that wild-type Piwi-bound sperm RNAs directly or indirectly trigger silencing of sequence-matching loci in the next-generation (zygote). This conclusion agrees with previous findings that maternal depletion of Piwi impacts heterochromatin formation in the offspring^28^. In reciprocal crosses, the paternal *aub*^*Het*^ mutation failed to trigger a similar piRNA dependent offspring transcriptional response (Fig. 3D) indicating a parent-of-origin directionality to the system. Thus, offspring transcription is modulated in a parent-specific manner by piRNA-pathway dosage.

**Fig. 3:**
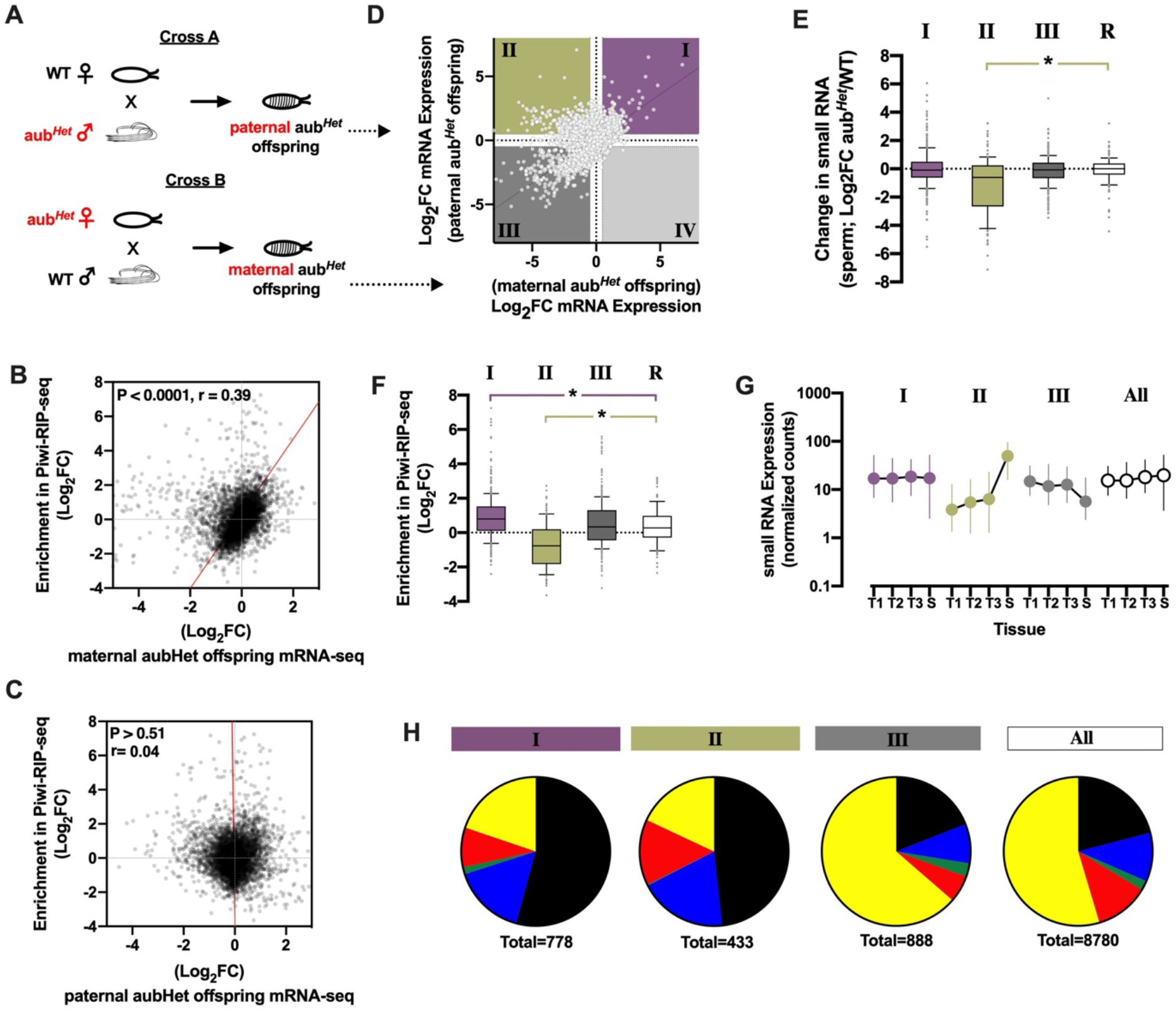
Paternal piRNAs reprogram offspring. A) crossing scheme; B) comparison of mRNA expression changes induced by paternal vs maternal *aub*^*Het*^, coloring and numbering is kept consistent throughout manuscript; C) correlation between Piwi pulldown enrichment and mRNA expression changes in maternal *aub*^*Het*^ offspring; D) correlation between Piwi pulldown enrichment and mRNA expression changes in paternal *aub*^*Het*^ offspring; E) Log_2_FC of sperm sRNAs in *aub*^*Het*^ vs WT males, each boxplot represents genes present in a different quadrants (Class) in D, random gene set in white F) Enrichment (Log_2_FC IP/Input) of sequence matching sRNAs in Piwi pulldown (Piwi-RIP-seq), each boxplot represents sequences matching genes present in a different quadrant (Class/color) from E; G) normalized counts in sRNA testis samples (as shown in Fig. 1) mapping to features in different quadrants in D, all genes in white; H) chromatin state annotations of Class I-III genes compared to all genes (right). Paternal *aub*^*Het*^ offspring n=5, WT offspring n=5, maternal *aub*^*Het*^ offspring n=2 with each replicate containing a pool of 20 embryos.

To gain further insights into the nature of parent-specific transcriptional changes in offspring, we compared the changes triggered in maternal-*aub*^*Het*^ offspring to those of paternal-*aub*^*Het*^ offspring, and identified three classes of intergenerationally-responsive transcripts and associated sRNAs: **Class I** transcripts (Fig. 3B **Purple**; n=1046) were upregulated in offspring of both maternal and paternal *aub*^*Het*^ crosses (Log_2_FC > 0.5). Class I transcript sequence matching sRNAs were enriched in our Piwi-RIP-seq indicating these transcripts are ‘targets’ of bona fide Piwi-bound sperm piRNAs (Fig. 3F). Class I transcripts were enriched for transposons (Supplemental Fig. 2A) which is in agreement with the canonical role of piRNAs in transposon silencing. Together, these data indicate that Class I genes in the embryo are targets of paternal piRNAs and their repression in the offspring requires zygotic aub activity. They identify a novel intergenerational sequence-specific gene regulatory axis.

**Class II** transcripts (n=494), by contrast, were upregulated *only* in offspring of paternal *aub*^*Het*^ crosses (Fig. 3B, **Green**; paternal Log_2_FC > 0.5; maternal Log_2_FC < −0.5). sRNAs mapping to Class II transcripts were depleted in the Piwi-IP (Fig. 3E) and, interestingly, were highly and specifically expressed in sperm (Fig. 3G). Further experiments revealed that sRNAs mapping to Class II transcripts were sensitive to both *aub* and *piwi* dosage in the male germline (reduced expression in *aub*^*Het*^ (Fig. 3E) and in *piwi*^*Het*^ (Supplemental Fig. 2B) sperm sRNAseq). These data identify a novel subclass of highly expressed sRNAs in sperm whose abundance is sensitive to aub and piwi dosage. Thus, Class II genes are intergenerationally regulated transcripts responsive to paternally inherited, piRNA-pathway sensitive sRNAs.

**Class III** transcripts (n=1036) were down regulated in offspring of both maternal and paternal *aub*^*Het*^ crosses (Fig. 3B, **Dark Grey**; Log_2_FC < −0.5). The Class III genes had no detectable signatures associated with transposon content (Supplemental Fig. 2A), Piwi-binding (Fig. 3F), or piRNA-pathway sensitivity (Fig. 3E and Supplemental Fig. 2B). There were only 20 **Class IV** transcripts that were specifically upregulated in maternal crosses (and only six with sequence-matching sperm sRNA reads, Fig. 3B, **Light Grey;** paternal Log_2_FC < −0.5; maternal Log_2_FC > 0.5). Importantly, to rule out potentially confounding influences of the arbitrary expression change cutoffs used above we repeated the analysis using a threshold free stratified *R*ank-*R*ank *H*ypergeometric *O*verrepresentation (RRHO) approach ^29^(Right side of Fig. S2). RRHO validated the interpretations above and the existence of Class I, II, and III transcripts (Fig. S2C) and their associated sRNA signatures (Fig. S2 D-F).

Gene set over-representation analysis showed enrichment in signaling and neuronal pathways (Fig. S2G top) for Class I (purple) genes. Class II (green) genes were enriched for metabolic, hydrolase activity and neuronal pathways (Fig. S2 middle), and Class III genes (dark gray) were enriched for cell cycle pathways (Fig. S2G bottom) genes. Consistent with the enrichments observed in our sperm Piwi-RIP-seq data (Fig. 3F), Class I genes were mostly positioned in “Black” Lamin / H1- and “Blue” Polycomb-associated repressive chromatin annotations (Fig. 3H, Class I). Interestingly, while not enriched in the IP, Class II genes also showed similar chromatin state association (Fig. 3H, Class II). Both of these annotations have been suggested to include enrichments for developmentally naive chromatin compartments and the data are therefore coherent with a mechanism for modulation of early embryo development. No such enrichment was found for Class III genes (Fig. 3H, Class III). These data are consistent with a mechanism whereby Class I and Class II *sRNAs* promote targeted, chromatin state specific silencing of offspring gene transcription. Thus, *paternal Piwi-bound sRNAs* and *paternal PIWI-sensitive sRNAs*, respectively, are necessary for next-generation repression of Class I & II genes.

### The piRNA-pathway is required for paternal inheritance of metabolic state

We previously identified similar chromatin-state and metabolic pathway signatures in offspring of a paternal diet-induced model of *I*nter*G*enerational *M*etabolic *R*eprogramming (IGMR)^30^. In that model (depicted in Fig. 4A), a two-day dietary sugar intervention in fathers before mating leads to obesity in adult offspring (∼10% increase in adult fat content), and in embryos, to transcriptional changes reminiscent of chromatin silencer deficiency. Guided by the similarities we tested for evidence of piRNA-pathway dependency in IGMR. We compared expression of the different Classes identified above (Fig. 3) to datasets from IGMR offspring. Intriguingly, while Class I genes showed no evidence of regulation in IGMR, Class II (piRNA-pathway dosage sensitive) genes showed significant upregulation in offspring of high sugar fed fathers (green; Fig. 4B), including strong enrichment of the most upregulated IGMR genes (upper bulge in the green violin; rank analysis, Fig. 4C). Embryonic transcription of Class II genes is thus sensitive to paternal dietary sugar; from a different point-of-view, diet-triggered intergenerational responses mimic those induced by aub and piwi heterozygosity in the male germline (paternal-*aub*^*Het*^ and *-piwi*^*Het*^). Signed rank correlation (Fig. 4D) and RRHO (Fig. S3A) analysis validated these findings genome-wide. Comparing IGMR and maternal-*aub*^*Het*^ -induced intergenerational responses did not show the same trend (Fig. 4D and S3B). The converse analysis, examining the most up- and down-regulated IGMR genes, confirmed these findings: Genes most upregulated in high sugar sired embryos showed the same signatures as Class II genes described above: they were upregulated in paternal, but downregulated in maternal *aub*^*Het*^ crosses (Fig S3C), and sRNAs mapping to these genes were downregulated in *aub*^*Het*^ mutant sperm and depleted from the Piwi pulldown (Fig S3D). Thus, paternal dietary sugar and piRNA-pathway deficiency trigger comparable intergenerational transcriptional rewiring in offspring. This indicates that IGMR is dependent on the dosage of sperm piRNA-pathway-regulated sRNAs.

**Fig. 4:**
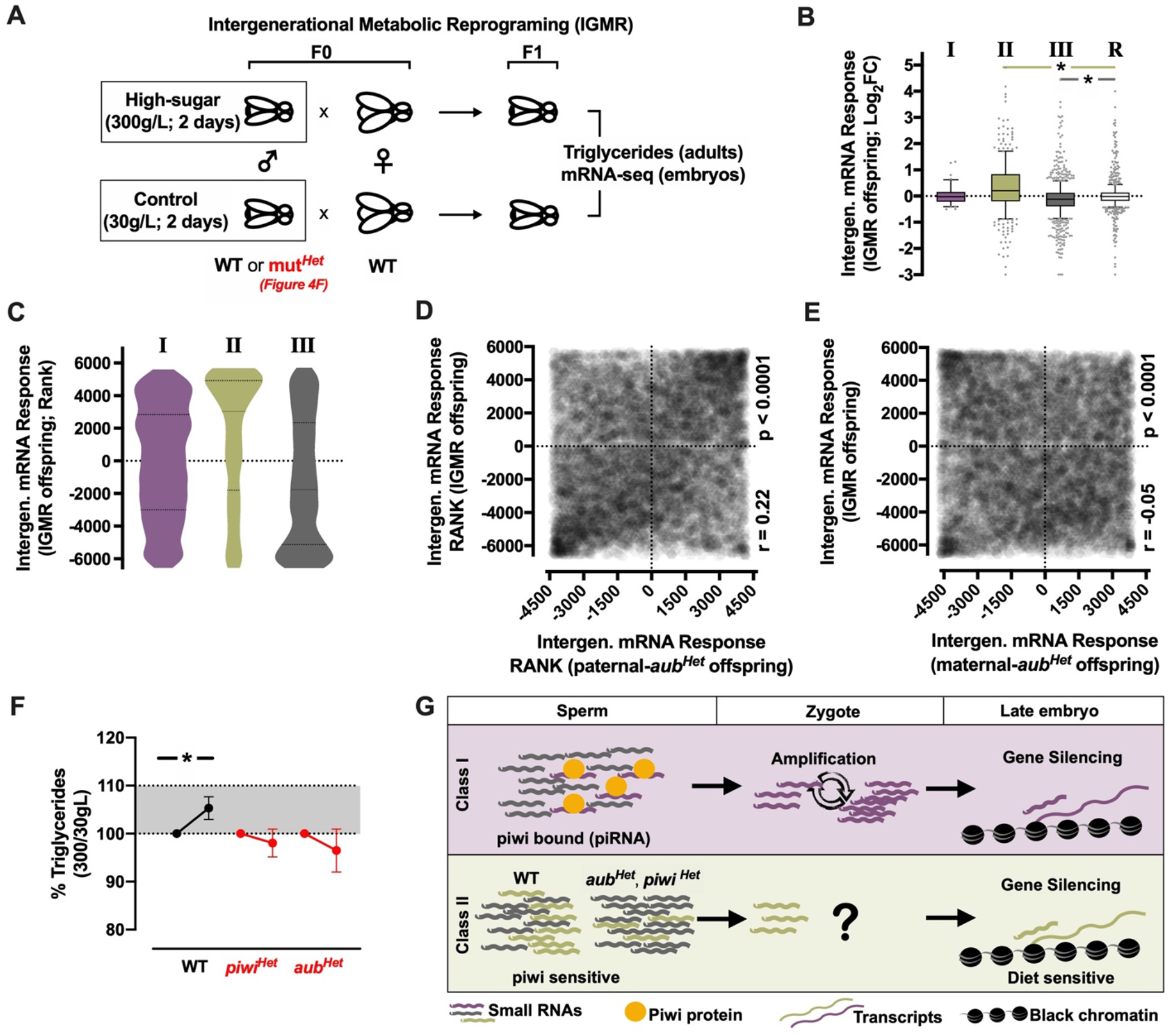
An intact piRNA-pathway is necessary for the intergenerational inheritance of a physiological phenotype. A) crossing scheme for paternal diet-induced intergenerational metabolic reprogramming (IGMR); B) mRNA-seq of IGMR offspring embryos, Log_2_FC control vs paternal high-sugar for different Classes as introduced in Fig. 3 B (R = random gene set); C) same as B but rank analysis instead of Log_2_FC; D) rank analysis comparing mRNA-seq of IGMR offspring embryos to paternal *aub*^*Het*^ offspring E) rank analysis comparing mRNA-seq of IGMR offspring embryos to maternal *aub*^*Het*^ offspring, p value and pearson r are depicted in D and E; F) whole body triglycerides of IGMR offspring as % of control-sugar matched value using wildtype (WT) or PIWI pathway heterozygous mutant (piwi, aub) fathers n=12-25 individual triglyceride assays of groups of 5 flies from at least 3 independent experiments; G) proposed model and summary for the data presented

If these piRNA sensitive genesets were causal with respect to intergenerational reprogramming, we reasoned that piRNA-pathway mutant fathers should fail to trigger an IGMR response. To this end, we tested whether piRNA-pathway heterozygote fathers (*piwi*^*Het*^ and *aub*^*Het*^) were capable of eliciting a full IGMR obesity response using our previously published intergenerational dietary model. We performed the intergenerational diet experiment using WT and *piwi*^*Het*^ and *aub*^*Het*^ fathers. Despite normal fertility and reproductive function in *piwi*^*Het*^ and *aub*^*Het*^ fathers, we found that both heterozygote lines failed to elicit increased triglyceride accumulation in the next generation (Fig. 4F). Thus, full piRNA-pathway dosage is necessary for paternal diet-induced intergenerational obesity (IGMR).

In summary, we describe a highly dynamic sRNA repertoire during *Drosophila* spermatogenesis. We find that sperm contains full-length piRNA-pathway proteins, and, using pulldown approaches, prove the existence of Piwi-bound piRNAs in sperm. Heterozygous mutations in piRNA-pathway member proteins lead to changes in sperm sRNAs indicating that mild pathway disruption is sufficient to alter the sperm sRNA load transferred to the zygote *and* to alter transcription in the next generation early life. Significant correlation between paternal Piwi bound (Class I) and Piwi sensitive (Class II) sRNAs and offspring gene transcription indicates that paternal sRNAs are involved in intergenerational inheritance by targeting sequence-matched genes for silencing in the next generation. Indeed, using paternal sugar triggered IGMR as a test case, we confirm this hypothesis and provide genetic evidence that a fully intact piRNA-pathway is necessary for intergenerational inheritance of paternal metabolic state (Summarized in Fig. 4G).

---

**Fig. S1:**
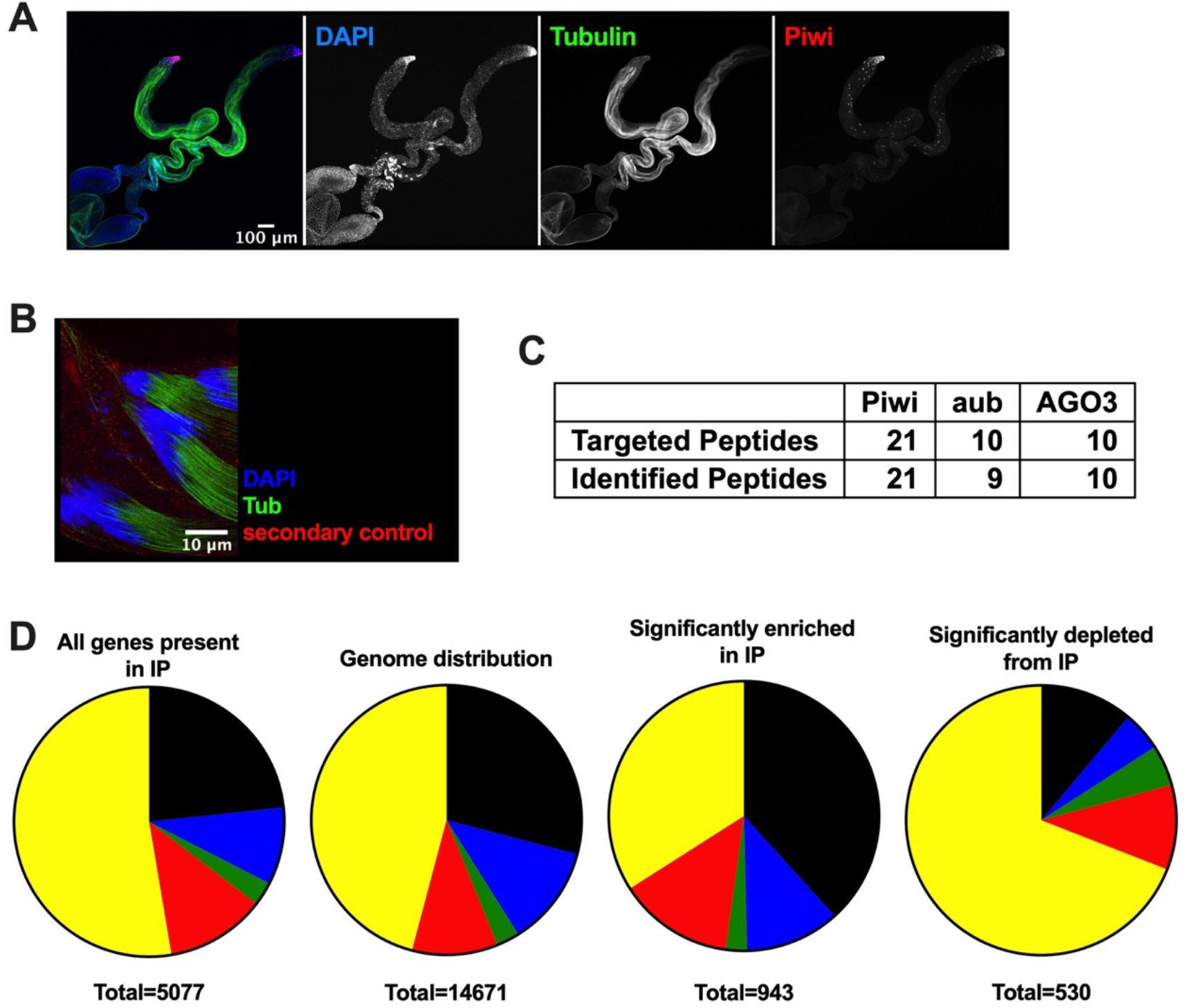
Presence of Piwi bound RNAs in mature sperm. A) immunofluorescence staining of testis with Piwi (sc-390946) and tubulin antibodies; B) secondary antibody control staining for Piwi (ab5207) staining in Fig. 2A; C) results of targeted mass spectrometry experiment identifying peptides in isolated sperm; D) pie chart representation of chromatin state distribution of reads mapping to features embedded in different chromatin states.

**Fig. S2:**
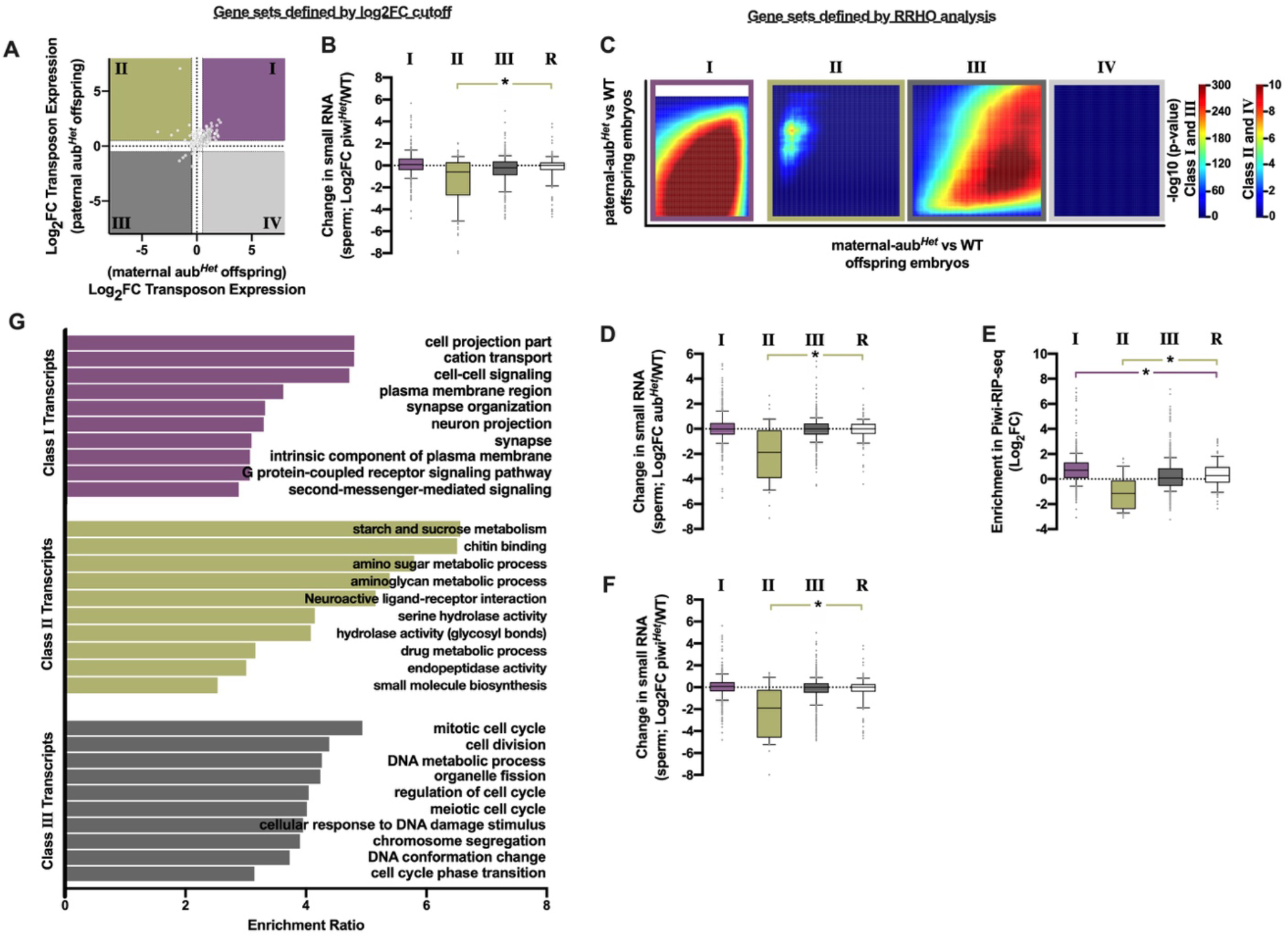
Paternal piRNAs reprogram offspring. A) comparison of transposon expression changes induced by paternal and maternal aub^*Het*^; B) Log_2_FC of sperm sRNAs piwi^*Het*^/WT, each boxplot represents genes present in a different quadrants (Class) from Fig. 3B, random gene set in white; C) Rank Rank Hypergeometric test of expression changes induced by paternal and maternal *aub*^*Het*^ confirms three Classes, different scales were applied to Class I/III and Class II/IV as depicted by the scale on the right; D) Log_2_FC of sperm sRNAs *aub*^*Het*^/WT, each boxplot represents genes present in a different quadrants from RRHO analysis in C, random gene set in white; E) Enrichment (Log_2_FC IP/Input) in piwi pulldown experiment, each boxplot represents genes present in a different quadrants from RRHO analysis in C, random gene set in white; F) Log_2_FC of sperm sRNAs *piwi*^*Het*^/WT, each boxplot represents genes present in a different quadrants from RRHO analysis in C, random gene set in white; G) over representation analysis of different Classes depicted in Fig. 3B using WebGestalt on gene ontology terms included in Biological Processes noRedundant

**Fig. S3:**
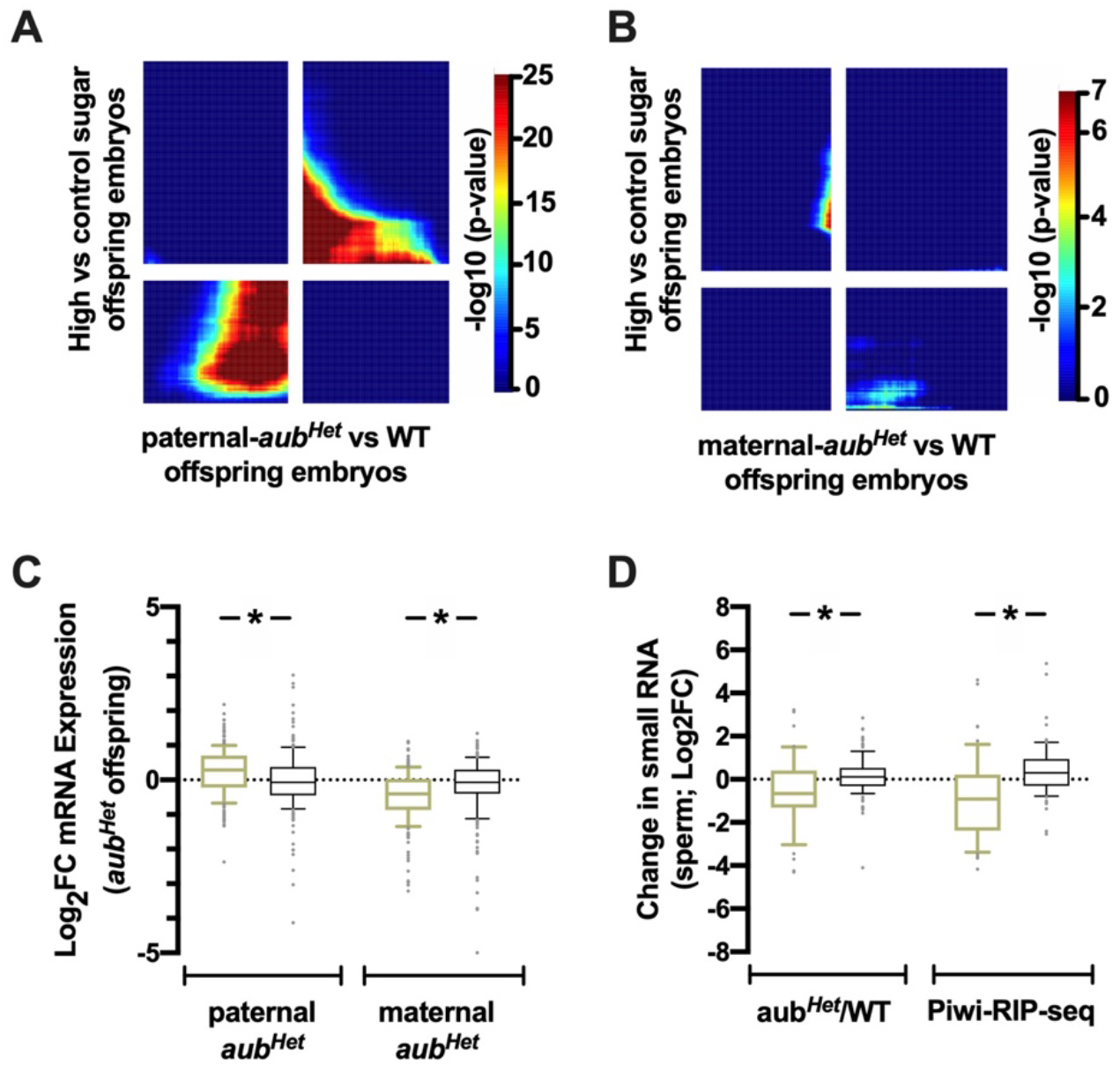
An intact piRNA-pathway is necessary for the intergenerational inheritance of a physiological phenotype. Rank Rank Hypergeometric test of expression changes in embryos induced by paternal diet intervention and A) paternal *aub*^*Het*^ B) maternal *aub*^*Het*^; C) Log_2_FC mRNA expression changes of the 300 most upregulated genes in IGMR offspring in paternal (green, left) and maternal (green, right) *aub*^*Het*^ offspring; D) Log_2_FC changes of sperm sRNA abundance in *aub*^*Het*^ sperm (green, left) and IP/Input (green, right), for those sRNAs mapping to the 300 most upregulated genes in IGMR offspring random gene set in black.

## Author Contributions

A.L., A.Ö. and J.A.P. conceived, designed and supervised the study. A.L. and J.A.P wrote the manuscript with feedback from all authors. A.L., L.E., A.G.M, O.L., M.S, U.K., M.I., L.Ö., M.G.L.R, E.B., performed fly work and dissections. U.K. perform library preparation for small RNA sequencing. L.E. and M.I. optimized and performed piwi RIP-seq. A.L. performed immunostaining. A.G.M. performed western blot analysis. T.R. prepared samples for mass spec. E.B. performed embryo mRNA sequencing. E.C., I.B., D.N., A.L., D.P. performed bioinformatic analysis. A.L. and T.V. supervised bioinformatic analysis.

## Supplementary Materials

### Materials and Methods

#### Fly Food

Standard food: Agar 12 g/l, yeast 18 g/l, soy flour 10 g/l, yellow cornmeal 80 g/l, molasses 22 g/l, malt extract 80 g/l, Nipagin 24 g/l, propionic acid 6,25 ml/l. Holidic diet was prepared according to ^32^.

#### Fly stocks

w1118: used as WT stock

aub^Qc42^ – BDSC #4968: used for genetic crosses

piwi^1^ – BDSC #43637: used for genetic crosses

T(2;3)TSTL, CyO: TM6B, Tb[1] – BDSC #42713: double balancer used for back-crossing

#### Fly Husbandry

Fly stocks were maintained on standard-diet at 25 °C on a 2-week generation cycle. To ensure a common parental larva density and epigenetic background, 0-4 days old flies were manually sorted to 15 males and 15 females per vial (or 45+45 in bottles) and allowed to lay eggs for 4 days. In practice, flies were crossed on Thursday, flipped out on Monday and virgins collected the following Monday. On Thursday flies were crossed again fulfilling a 2-week generation cycle.

#### Generation of flies for genetic crosses

To equalize the genetic background between mutant and WT strains we crossed the mutant lines over double balancer lines so chromosomes can be followed using a visible marker. These flies were then crossed to the WT strain and all chromosomes were replaced with WT chromosomes except the one carrying the mutation and the respective balancer. Flies from the above cross were crossed with WT and the resulting offspring without the balancer chromosome, was used for the for the paternal and maternal mut^*Het*^ crosses.

#### Testis, sperm sack and ovary dissection

The reproductive tract of separated male and female flies was dissected in a drop of TC-100 insect medium (Sigma) and connective tissue and other contaminating tissues were removed. Ovaries were transferred to a tube containing insect medium. Testes with attached seminal vesicles (sperm sacks) were transferred to a fresh drop of insect medium, where the sperm sack was separated from the rest of the testis. Testes were further separated into three parts and transferred to different tubes containing insect medium. Mature sperm was removed from the sperm sacks by puncturing them and spooling the sperm onto clean forceps and transferred to a tube containing insect medium. After a maximum of 30 min of dissection, collected tissues were snap frozen in liquid nitrogen. When enough samples were collected the tissues were pelleted, the supernatant was discarded, and the sperm pellet was re-suspended in 250µl of Trizol (Invitrogen).

#### Collection of staged *D*.*melanogaster* embryos

Virgin female flies were kept in egg-laying cages with apple-juice plates supplied with fresh yeast paste for three days. Female virgin flies were crossed to male virgin flies in egg-laying cages at 25 °C in the morning and apple juice plates were changed approximately every 30 min until 2pm. Subsequent apple juice plates were used for collection of embryos. To obtain stage 17 embryos pates were left on the cage for 2 h and incubated for 16 h at 25 °C. Embryos on apple juice plates were dechorionated using 50% PBS-TX (PBS-T containing 0.3 % Triton X-100) and 50 % bleach for ∼ 2 min. Dechorionated and detached embryos were collected and rinsed under a stream of water. Embryos were then staged according to their morphology to maximize homogeneity in the sample and immediately transferred to 250 µl of Trizol solution for RNA isolation.

#### RNA isolation for sRNA-seq

Samples prepared in Linkoping: Frozen samples were homogenized in Qiazol with 0.15 g 0.2 mm steel beads, Tissue Lyser 2 min 40 osc. RNA extraction was done using miRNeasy Micro kit (Qiagen, Venlo, the Netherlands) and performed according to the manufacturer instructions, RNA was eluted in 14 μl of water and stored at −70 °C until library preparation.

Samples prepared in Freiburg: Dried RNA were shipped on dry ice and kept in −70 °C. Dried and washed once with 80% ethanol and dried, dissolved in 20 ul water, extraction was done using miRNeasy Micro kit (Qiagen, Venlo, the Netherlands) and performed according to the manufacturer instructions. RNA was eluted in 14 μl of water and stored at −70 °C until library preparation. Bioanalyzer confirmed the quality of the RNA.

#### Library preparation for sRNA-seq

Library preparation was done with NEBNext Small RNA Library Prep Set for Illumina (New England Biolabs, Ipswich, MA) according to the manufacturer instructions with the following minor customizations. All testicle samples, but not sperm samples, were downscaled to half volume, using 3 μl of input RNA instead of 6 ul as recommended. In all steps the primers in the kit were diluted 1:3 prior to use. 2S rRNA was blocked by adding anti-sense oligos (5’-TAC AAC CCT CAA CCA TAT GTA GTC CAA GCA372 SpcC3 3’; 10 µM) and set to hybridized in the same step as NEBNext SR RT primer (pink). Amplification was made during 16 cycles and amplified libraries were cleaned using Agencourt AMPure XP (Beckman Coulter, Brea, CA) and size selected for 130 to 165 nt fragments on a pre-casted 6% polyacrylamide Novex TBE gel (Invitrogen, Waltham, MA). Gel extraction was done using Gel breaker tubes (IST Engineering, Milpitas, CA) in the buffer provided in the NEBNext kit. Disintegrated gels were incubated at 37 °C for 1 hour on a shaker, quickly frozen for 15 minutes at −80 °C, followed by another incubation for 1 hour. Any remaining gel debris was removed by Spin-X 0.45 μm centrifuge tubes (Corning Inc., Corning, NY) as recommended by the NEBnext protocol. The libraries were precipitated overnight at −80 °C by adding 1 μl of GlycoBlue (Invitrogen) as co-precipitant, 0.1 times the volume of Acetate 3M (pH 5.5), and 3 times the volume of 100% ethanol. Library concentrations were estimated using QuantiFluor ONE ds DNAsystem on a Quantus fluorometer (Promega, Madison, WI). Pooled libraries were sequenced on NextSeq 500 with NextSeq 500/550 High Output Kit version 2, 75 cycles (Illumina, San Diego, CA). All pooled libraries passed Illumina’s default quality control.

#### Immunostaining

Immunostaining of whole mount testis and dissected sperm was carried out according to^33^ in short: testes were dissected in insect medium (Sigma, TC-100) and all incubation and washing steps were carried out using home-made baskets in 96 well plates on an orbital shaker. Fixation was carried out in 4% formaldehyde (methanol-free) in PBST (PBS and 0.2% Triton X-100) for 10 min at room temperature. After three 5 min washing steps in PBST testes were permeabilized twice in 0.3% sodium-deoxycholate in PSTX for 30 min at room temperature. This was followed by three 5 min washing steps in PBSTX and 1h of blocking with 5% BSA in PBSTX at room temperature. Primary antibody incubation (anti-Piwi antibody ab5207 - 1:500 dilution; anti-acetylated α-tubulin (Lys40) antibody – 1:500, anti-Piwi antibody sc-390946 – 1:500 dilution) was carried out in PBSTX+3% BSA at 4 °C overnight followed by one 20min wash with 300 mM NaCl in PBSTX and three 5 min washes with PBSTX at room temperature. Secondary anti-Mouse Alexa Fluor 488 (1:1000; Molecular Probes) and anti-rabbit Alexa Fluor 555 (1:1000; Molecular Probes), incubation lasted for 6h in PBSTX + 3% BSA at 4 °C samples were washed with 300 mM NaCl in PBSTX for 20 min at room temperature, all steps were repeated for the second primary and secondary antibodies and samples were mounted with Vectashield with DAPI (Vector Labs). Confocal images were taken with a Zeiss LSM510 confocal scanning microscope with a C-Apochromat × 63, 1.4 NA oil immersion objective, using the diode 405 nm, the argon 488 nm, the helium–neon 543 nm laser for excitation of DAPI, Alexa Fluor® −488, −555, respectively.

#### Triglyceride Determination

Groups of five flies (7-12 days old males) were crushed thoroughly in 100 µl RIPA buffer, sonicated and the homogenates were used for 96-well based colorimetric determination of triglycerides (GPO Trinder, Sigma). Before absorbance measurement, plates were centrifuged, and supernatants transferred to a new plate.

#### Mass Spectrometry (MS/MS) analysis

Sperm sack and pure sperm samples were analyzed by nanoLC-MS. Cells were lysed with lysis Buffer (4% SDS, 100mM DTT,50 mM Tris-HCl buffer pH 7.5, plus protease inhibitors (Roche)). Then the extract was heated to 90°C for 3 min followed by residual chromatin shearing in Bioruptor. Before MS-MS analysis samples were analyzed by SDS-PAGE followed by silver staining (Thermofisher).

#### Western blotting

For protein extraction tissue were lysate adding 50 ml of Laemmli sample buffer 2x, heated at 95°C for 2 min and sheared in Biorupter (30sec/hard). Protein lysates were loaded on NUPAGE 4-12% precast gel (Life Technologies) in NUPAGE 1xMOPS buffer (Invitrogen). PageRuler Plus Prestain Protein Ladder (Thermo Scientific) was used to indicate protein size. Proteins were subsequently transferred to PVDF membranes. Before blocking, membranes were stained with Ponceau solution (Sigma) and images were recorded. Membranes were blocked in phosphate buffered saline plus 0,05% Tween 20 (PBST) and 5% BSA for 1 h at room temperature. Membranes were incubated with primary Anti-Piwi antibody (ab5207) at 1:500 dilution overnight at 4 °C, washed in PBST, and incubated with horseradish peroxidase (HRP) coupled secondary antibodies (Anti-rabbit IgG, HRP-linked Antibody #7074) in the washing buffer with 1% skimmed milk in PBST for 1 h at room temperature. Membranes were developed using SuperSignal™ West Femto Maximum Sensitivity Substrate (Thermofisher).

#### Ribonucleoprotein Immunoprecipitation (RIP)

All steps were performed on ice or at 4 °C unless indicated otherwise. Samples with magnetic beads were placed on magnetic rack for 30sec to ensure beads were pelleted. Dissected tissues were thawed on ice and resuspended in PBS-T (0.05% Tween 20) supplemented with cOmplete, mini EDTA-free protease inhibitor cocktail (Roche Diagnostics, GmbH, Germany) and RNase inhibitor. Samples were transferred to a 60mm cell culture dish (on ice) and placed in a Bio-LinkTMBLX 365 UV-crosslinker. Samples were crosslinked with UV light at 400 mJ/cm2. Samples were then transferred to a fresh 1.5 ml Eppendorf tube. 60 mm dish was then washed once with 500 μl PBS-T and the wash was added to sample. Samples were then centrifuged 8000 x g at 4 °C for 5 min. Supernatant was carefully discarded and samples were resuspended in 250 μl-500 μl RIPA buffer (50mM Tris-HCl pH 7.5, 150 mM NaCl, 1% Triton-X 100, 0.1% SDS, 0.1% Na-deoxycholate, 1mM EDTA, 1x cOmplete, mini EDTA-free protease inhibitor cocktail, RNase inhibitor) and transferred to a Wheaton dounce tissue homogenizer chilled on ice. Samples were homogenized 10x with a loose pestle and then 50x with a tight pestle to ensure complete tissue homogenization. The homogenized samples were transferred to a fresh 1.5 ml Eppendorf tube and kept on ice. The homogenizer was rinsed with 250 μl −500 μl IP dilution buffer (50 mM Tris-HCl pH 7.5, 150 mM NaCl, 1x cOmplete, mini EDTA-free protease inhibitor cocktail, RNase inhibitor) so as to dilute RIPA buffer 1:1 and the wash solution transferred to same tube as homogenized sample. A 5% aliquot was taken of the lysate to be used as either input for IP-western blot experiments or input for small RNA sequencing experiment. Sample stored at −20 °C.

Meanwhile, Diagenode CHiP-kit protein A magnetic beads were washed before use with 500ul PBS-T and then 500 μl IP buffer (1:1 RIPA buffer and IP dilution buffer). Samples were pre-cleared with pre-washed 25 μl Diagenode CHiP-kit protein A magnetic beads at 4 C with end-to-end rotation for 2 hrs. A second batch of pre-washed Diagenode CHiP-kit protein A magnetic beads were pre-loaded with rabbit-@-PIWI (ab5207) or IP control rabbit IgG: either 4 μg antibody for 25 μl bead slurry for IP-Western blot experiments or with 8 μg antibody for 50 μl bead slurry for IP-targeted proteomics experiment/smallRNA library preparation in 500 μl IP buffer (1:1 RIPA buffer and IP dilution buffer). The bead/antibody solution was incubated at 4 °C for 4-6 hours with end-to-end rotation. Pre-cleared samples were then pelleted on a magnetic rack and supernatant transferred to the preloaded beads (following the disposal of the supernatant). Antibody/preloaded beads/pre-cleared lysate mixture was incubated at 4 °C overnight with end-to-end rotation. The following day, samples were washed 7 x 1 ml wash buffer (50 mM Tris-HCl pH 7.5, 150 mM NaCl, 2 mM MgCl2, 10% glycerol, 1% Empigen, RNase inhibitor) ^34^. Each wash was carefully removed using vacuum suction. Samples used for IP-western blot or IP-targeted proteomics experiments were eluted with 30 μl Bolt LDS sample buffer (Novex, Life Technologies) supplemented with Bolt sample reducing agent (Novex, Life Technologies) at 95 °C, 1000 rpm for 5min then chilled on ice. Samples were stored at −20 °C until needed. If RNA was needed following PIWI immunoprecipitation, beads were incubated with 100 μl TRI Reagent® for 10 min at room temperature. Input samples were incubated with 10 volumes of TRI Reagent® for 10 min at room temperature.

#### RNA isolation for small RNA library preparation

Tubes containing phase lock gel were prepared by the filling the lids of 0.2 ml PCR tubes with phase lock gel. The tubes were then quickly spun down. Following incubation with TRI Reagent®, the beads were pelleted using a magnetic rack and the supernatant was transferred to a tube containing phase lock gel. Subsequently, 20 μl chloroform was added, the tubes mixed by shaking and then centrifuged at 3100 x g for 10 min. The upper phase was transferred to a fresh tube containing 50 μl isopropanol and 15 μg GlycoBlueTM, mixed by shaking and then incubated at − 20 °C overnight. The samples were then centrifuged at 3100 x g for 10min at 4 °C. The RNA pellet was washed with 100 μl 75% ice cold ethanol, centrifuged at 3100 x g for 10min at 4 °C. Supernatant was carefully decanted off samples and the pellet air dried. The pellet was then resuspended in 4 μl nuclease free water and stored at −80 °C. 5% of lysate used for PIWI IP (25 μl out of 500 μl lysate) was reserved to be used as input for small RNA sequencing experiment. These samples were incubated with 250 μl TRI Reagent® for 10min at room temperature. Subsequently, 50 μl chloroform was added, the tubes mixed by shaking and then centrifuged at 3100 x g for 10min. The upper phase was transferred to a fresh tube containing 125 μl isopropanol and 15 μg GlycoBlueTM, mixed by shaking and then incubated at −20 °C overnight. The samples were then centrifuged at 3100 x g for 10min at 4 °C. The RNA pellet was washed 1x with 250 μl 75% ice cold ethanol, centrifuged at 3100 x g for 10min at 4 °C. Supernatant was carefully decanted off samples and the pellet air dried. The pellet was then resuspended in 6.5 μl nuclease free water and stored at −80 °C.

IP and Input samples processed the same way from now on: The quality and concentration of the RNA samples was analyzed by Agilent Small RNA kit on the Agilent 2100 Bioanalyzer system and NanoDropTM. The RNA samples were then used to make small RNA libraries using the NEBNext Small RNA Library Prep Kit for Illumina (E7330) according to the manufacturer’s instructions. Ligation of the 3’SR adapter was performed at 16 °C for 18 hrs. This longer incubation at a reduced temperature increases ligation efficiency of methylated RNAs such as piRNAs. Since *Drosophila* RNA is rich in 2S rRNA, a 2S blocking oligo was also used at a final concentration of 0.1 μM to exclude 2S RNA from any downstream reaction. 12 PCR cycles were used to amplify the libraries.

#### QC and read mapping

Small RNA reads were trimmed for sequencing adapters using Trim Galore v0.5.0 (https://github.com/FelixKrueger/TrimGalore) in conjunction with Cutadapt v1.11^35^. Low-quality (Q<20) bases were also trimmed from the ends of reads, and reads were discarded if their length was then less than 20 nucleotides following trimming.

Trimmed reads greater than 20 nucleotides in length were then mapped to dm6 genome (FlyBase BDGP6.22 release) using the short-read aligner bowtie v1.2.3 ^36^, allowing for one mismatch. If there were multiple alignments, then only the highest quality alignment was retained (bowtie options -M 1 --best --strata). Alignment files were sorted and indexed using SAMtools v1.8.

#### Hierarchical Feature Counting

Annotation data was downloaded from FlyBase, release BDGP6.22. The annotations were stratified into 14 ordered feature categories: rRNA, tRNA, snRNA, snoRNA, pre miRNA, simple repeats, complex repeats, Piwi RNA, 5’ UTR, 3’ UTR, protein coding exon, pseudogene, ncRNA, and mitochondrial genome. The Piwi RNA cluster (n=114) annotations were from piPipes. Next, using featureCounts (subread v2.0.0)^37^, mapped reads were counted for each of the 13 categories independently. A minimum overlap of 15 nucleotides was required, with the fractional counts option was on. This counting was strand-specific and was done for both forward (sense) and reverse (antisense) strands independently. Finally, using a custom script (https://github.com/vari-bbc/Piwi_RNA_pipeline), we did a hierarchical counting for the 13 categories. That is, for any read annotated to multiple features, it was counted only for the feature highest in the category. So, reads mapped to a Piwi RNA cluster within an exon were counted towards the Piwi RNA cluster not the exonic feature. This approach assured that reads were assigned correctly, and ambiguities from reads mapping to rRNA and other small RNA features were not included as Piwi RNAs. This resulted in a counts file for sense and antisense features.

#### Differential Expression Analysis

The sense and antisense counts tables were imported into R v3.6.0. Features with counts of less than 10 raw counts in less than 2 samples were removed. In addition, two complex repeat features (LSU-rRNA_Dme and SSU-rRNA_Dme) were also removed. Following filtering, individual contrasts for differential feature expression were done using edgeR v3.28.0^38^. For all contrasts, a general linear model was fit using appropriate covariates that varied depending on the contrast, but included – where appropriate – treatment, tissue, knock, knock side, and genotype. Statistical significance was assessed using a quasi-likelihood F test with multiple testing correction performed with the Benjamini-Hochberg procedure.

#### GSEA

The genes comprising each biotype were used as gene sets to test for enrichment of certain biotypes in certain phenotypes. Enrichment testing was done with GSEA as implemented in the clusterProfiler v3.14.3 function ‘GSEA.’ Then thousand phenotype permutations were run ^39^. The biotype annotation was from UCSC for dm6.

#### Overrepresentation Analysis (ORA)

ORA for RIP enriched transcripts was performed using WebGestalt ^31^ with the following parameters for the enrichment analysis: Minimum number of IDs in the category: 5; Maximum number of IDs in the category: 500; FDR Method: BH; Significance Level: FDR < 0.05

